# Syndecan Regulates Cellular Morphogenesis in Cooperation with the Netrin Guidance Pathway and Rho-family GTPases

**DOI:** 10.1101/2022.01.13.476274

**Authors:** Raphaël Dima, Marianne Bah Tahé, Yann A. Chabi, Lise Rivollet, Anthony F. Arena, Alexandra M. Socovich, Daniel Shaye, Claire Y. Bénard

**Author notes:** Equal contribution (Raphaël Dima, Marianne Bah Tahé). Corresponding authors: Claire Bénard, Université du Québec à Montréal, Department of Biological Sciences, 141 President Kennedy Avenue, Montréal, QC H2X 1Y4, Canada, extension -6192, Daniel Shaye, University of Illinois at Chicago – College of Medicine, Department of Physiology and Biophysics (MC901), 835 S. Wolcott Ave, MSB Room E202, Chicago, IL 60612, USA.

## Abstract

The establishment of complex cell shapes is essential for specific cellular functions, and thus critical in animal development and physiology. Heparan sulfate proteoglycans (HSPGs) are conserved glycoproteins that regulate interactions between extracellular signals and their receptors, to orchestrate morphogenetic events and elicit cellular responses. Although HSPG-regulated pathways have been implicated in regulating the guidance of neuronal migrations, whether HSPGs regulate earlier aspects of cellular development that dictate cell shape remains unknown. HSPGs consist of a protein core (e.g., Syndecan, Perlecan, Glypican, etc.) with attached heparan sulfate (HS) glycosaminoglycan chains, which are synthesized by glycosyltransferases of the exostosin family. Using mutations in the two *C. elegans* HS glycosyltransferases genes, *rib-1* and *rib-2*, we reveal that HSPGs control the number of cellular projections in the epithelial excretory canal cell, which can form more than its normal four canals in these mutants. We identify SDN-1/Syndecan as the key HSPG that regulates the number of excretory canal cell projections in a cell-autonomous manner. We also find that Syndecan and guidance receptors for Netrin function in the same pathway to restrict the number of cellular projections. Furthermore, we show that the formation of extra projections in the absence of Syndecan requires the conserved Rho-family GTPases CED-10/Rac and MIG-2/RhoG. Our findings not only contribute to understanding the roles of conserved HSPGs in cellular morphogenetic events, but also reveal the existence of an HSPG-regulated system operating to guarantee that a precise number of cellular projections is established during cell development. Given the evolutionary conservation of developmental mechanisms and the molecules implicated, this work provides information relevant to understanding the cellular and molecular bases of the development of precise cellular morphologies in varied cell types across animals.

## INTRODUCTION

The establishment of complex cell shapes is essential for specific cellular functions, and thus critical is in animal development and physiology. Certain cells possess a specialized morphology that allow distinct functions such as the flow of information along a neuron, or the osmoregulation carried out by the tubular cells of the excretory system. Whereas the mechanisms driving the outgrowth and guidance of neuronal projections have been uncovered and described over the past two decades, how the *number* of cellular projections is controlled during development remains poorly understood.

Cellular morphogenesis relies on mechanisms intrinsic and extrinsic to cells^1,2^, which bring about asymmetries within the developing cell and establish dynamic motile structures underlying oriented growth. In developing neurons, for instance, growth cones bear multiple receptors for extracellular guidance cues such as UNC-6/Netrin^3^. UNC-6/Netrin is a molecule with a key role in modulating the outgrowth and guidance of the extending growth cone, as well as of other polarized cells^4^. The UNC-6/Netrin signal can act as a growth cone attractant by interaction with a homodimer of its receptor UNC-40/DCC^4–6^, or as a repellent by interaction with an heterodimer of UNC-40/DCC and UNC-5/UNC5 receptors^7^ inducing destabilization of the cytoskeleton in the growth cone protrusions^8,9^. The UNC-6/Netrin guidance pathway is also involved cell symmetry breakage and the definition of the axon formation site, prior to its roles in the guidance of established cellular projections^2^.

Receptors of guidance cues recruit and activate intracellular signals to induce cell responses, which often involve cytoskeleton rearrangements. MIG-2/RHOG and CED-10/Rac are two small Rho-GTPases members of the Rac-like GTPases subfamily that function in the UNC-6/Netrin guidance pathway, to regulate the growth cone’s cytoskeleton dynamics^10^. MIG-2/RHOG and CED-10/Rac are also required for proper extension outgrowth of other cells such as those of the excretory canal cell in *C*. *elegans*^11^. Besides their role in guidance, these regulators also modulate the cytoskeleton during the earliest phase of cell extension or axonal specification^12^.

Heparan sulfate proteoglycans (HSPGs) are conserved glycoproteins that regulate interactions between extracellular signals and their receptors, to elicit cellular responses and orchestrate morphogenesis. They function as co-receptors, modulating the interaction of morphogens and guidance cues with their receptor, in numerous developmental pathways such as that of netrin^13,14^, among others^15^. HSPGs are composed of a core protein to which heparan sulphate (HS) glycosaminoglycan chains are attached. HS chains are synthesized by exostosin glycosyltransferases. The HSPGs core proteins^13,15^ as well as their HS chains can both play roles in the development of neurons and other cells with complex morphologies^15–17^, and the chemical modification of HS chains increases their specificity of function^15,18^.

Here, we uncover a mechanism that regulates the *number* of cellular extensions established by a cell during development. Using the excretory cell of *C*. *elegans*, we reveal that HSPGs, and more specifically, SDN-1/Syndecan is the key HSPG at the center of this mechanism. We find SDN-1 functions in a cell-autonomous manner together with the Netrin guidance receptors to restrict the number of cellular projections. Furthermore, we show that the formation of extra cellular projections in the absence of Syndecan requires the conserved Rho-family GTPases CED-10/Rac and MIG-2/RhoG. Together, our findings contribute to understanding the roles of conserved HSPGs in cellular morphogenetic events, in mechanism regulating the development of precise cellular morphologies.

## MATERIALS AND METHODS

### Nematode strains and genetics

Nematode cultures were maintained in an incubator at 20 °C (unless otherwise noted) on NGM plates seeded with *Escherichia coli* OP50 bacteria as described^19^. Alleles used in this study are listed in Table S3. Strains were constructed using standard genetic procedures and are listed in Table S2. Genotypes were confirmed by genotyping PCR or by sequencing when needed. Primers used to build strains are listed in Table S1. All the mutant alleles and reporter strains were outcrossed with the Bristol N2 wild-type strain at least 3 times prior to use for analysis or strain building.

### Microinjections and transgenic animals

Transgenic animals were generated by standard microinjection techniques^20^. Each plasmid was injected at 0.2 ng/μL (pCB420), 1 ng/μL (pCB427), 5 ng/μL (pCB436), 10ng/μL (pCB423) or 25 ng/μL (pCB242, pCB265, pCB425) as indicated in Table S5, with the co-injection marker P*unc*-*122::rfp* (coelomocytes in red) at a concentration of 50 ng/μL. pBSK+ was used to increase the total DNA concentration to 200 ng/μL.

### Microscopy and excretory canal cell observations

Worms were grown in an incubator at 20 °C for at least 3 generations prior to analysis. Worms were anesthetized with 75 mM NaN3 between a coverslip and a 5% agarose pad. Observations were made using a ×40 or ×63 objective on a Zeiss Axio Imager (M2) fluorescence microscope. Images of representative phenotypes were captured using the Zeiss AxioCam camera and the Zen software. Images of embryos were deconvoluted using the software *AutoQuant* X3 for better resolution.

The excretory canal cell and its canals were examined in L4 stage larvae, unless otherwise specified, using the transgenic reporters *arIs164* (P*glt*-*3::venus*) or *bgIs312* (P*pes*-*6::gfp*). The excretory canal cell is located near and ventral to the terminal bulb of the pharynx in the head and bears four lateral canals in the wildtype: two anterior canals located along each side of the animal’s head (one on the left and one on the right side), and two posterior canals located along the length of the animal’s body (one on the left side of the animal and one on the right side). An animal was considered to have supernumerary canals only when more than four canals had grown directly out of the excretory canal cell *soma*, regardless of the canal’s position in the animal’s body (left, right, dorsal, ventral, anterior or posterior). Only canals that (1) stem from the excretory cell *soma* and (2) are longer than the diameter of the excretory cell soma (∼ 25 μm in length) were counted as canals. Thus, branches that had extend along the length of a canal were *not* counted as an extra canal, as they do not arise from the excretory canal cell soma. Other types of defects were observed during our analyses such as missing canals, misguided canals, and shorter canals, as described elsewhere^4,21–23^, which were not counted as defective in this study as we focused on the presence of *supernumerary* canals.

### Rescue and expression constructs

Numbers in square brackets below refer to position along the cDNA.

P*mec*-*7::sdn-1* cDNA (pCB242): The cDNA of *sdn*-*1* was amplified from yk139f3 and ligated into a P*mec*-*7* containing pPD96.41 vector using enzymes *Xma*I – *XhoI* as described previously^13^.

P*glt*-*3::sdn-1* cDNA (pCB420): Vector pCB242 (P*mec*-*7::sdn-1* cDNA) was digested with *Sph*I [11] and *Xma*I [888] to release P*mec*-*7* and ligated with insert P*glt*-*3* [303 bp] amplified from vector pDS629 (P*glt-3::zif-1*) using primers oCBQc48 (GCTTGCATGCTTTCGAATCG) and oCBQc49 (CATGATCCCGGGTATGGATCCGGTACCTCC) to add on *Xma*I site, and then digested with *Sph*I [9] and *Xma*I [292].

P*grd-10*::*sdn*-*1* cDNA (pCB265): The cDNA of *sdn*-*1* was amplified with primers oCB1032 and oCB1033, digested with *Eco*RI and *Age*I and ligated into *grd-10*::GFP containing pPD95.75 backbone vector with GFP replaced by the cDNA of *sdn*-*1* with *Eco*RI and *Age*I.

P*rgef-1::sdn-1* cDNA (pCB425): Vector pCB199 (P*rgef-1::rib-1* cDNA) was digested with *Xma*I [3480] and *Apa*I [5441] to release the cDNA of *rib*-*1* and the *unc-54* 3’UTR and ligated with the fragment containing the *sdn*-*1* cDNA and *unc-54* 3’UTR amplified from pCB242 (P*mec*-*7::sdn-1* cDNA) digested with *Xma*I [888] and *Apa*I [2734].

P*myo-3::sdn-1* cDNA (pCB423): Vector pCB332 (P*myo-3::lon-2* cDNA) was digested with *Xba*I [2414] and *Sac*I [3998] to release the cDNA of *lon-2* and ligated with the fragment containing the cDNA of *sdn*-*1* amplified from pCB420 (P*glt-3::sdn-1* cDNA) using primers oCB1715 (CATGATTCTAGACCATACCCGGGATGATTCTG) and oCB1716 (AAGATCTCGGGAGCTCCTC) to add on *Xba*I site and then digested with *Xba*I [7] and *Sac*I [901].

P*rab-3::sdn-1* cDNA (pCB436): Vector pCB428 (P*rab-3::sax-7* cDNA) digested with *Xma*I [1528] and *Xho*I [5091] to release the cDNA of *sax-7* and ligated with the fragment containing the cDNA of *sdn*-*1* amplified from pCB420 (P*glt-3::sdn-1* cDNA) [4737 bp] digested with *Xma*I [294] and *Xho*I [1167].

P*myo-2::sdn-1* cDNA (pCB427): Vector pCB242 (P*mec*-*7::sdn-1* cDNA) was digested with *Xba*I [25] and *Xma*I [888] to release P*mec*-*7* promoter and ligate with insert P*myo-2* [863 bp] amplified from pCB230 (P*myo-2::rib-2* cDNA) digested with *Xba*I [25] and *Age*I [1309].

All inserts of finalized clones were verified either by sequencing for cloned PCR amplicons, or by restriction digests and PCR amplification for fragments released from an already sequenced vector.

## RESULTS

### The HSPG SDN-1/Syndecan limits the number of excretory canal cell projections

The excretory canal cell, which ensures osmoregulation in *C*. *elegans*, is an epithelial tubular cell whose soma is located in the head of the animal, and from which four projections or canals extend along the body sides, two anteriorly and two posteriorly (**Fig. 1A**, top panel). During previous analyses, we noticed excretory canal cell abnormalities in mutants affecting the heparan sulphate (HS) chains biosynthesis^16^. HS chain biosynthesis is catalyzed by a co-polymerase composed of two exostosin glycolsyltransfereases RIB-1/EXT1 and RIB-2/EXTL3 in *C*. *elegans* and mammals alike (**Fig. 1B**). In hypomorphic viable mutants *rib*-*1(qm32)* and *rib*-*2(qm46)*, the guidance of excretory canal cell projections, neurons, axons, and the distal tip cell, is defective^16^. The outgrowth per se of the excretory canal cell projections can also be affected in these mutants, as the excretory canals can be short (**Fig. 1A**). These mutants also frequently exhibit a striking and novel type of defect, where animals can have an excessive *number* of excretory canals, compared to the wild type. We characterized this defect using a reporter to visualize the excretory cell (P*pes*-*6::gfp*) and found that ∼ 27% of the *rib*-*1* and *rib*-*2* mutants develop supernumerary canals, ranging from five to eight excretory canals, compared to the four projections invariably observed in the wild type (**Fig. 1C**). Supernumerary excretory canals as long as the normal excretory canals, or shorter, located anteriorly or posteriorly from the soma of the excretory canal cell, or even in a ventral or lateral position were observed. Excretory canal cell projections or canals that grew from the *soma* of the excretory canal cell and that exceeded the set of four normally present in the wild type, are henceforth called “extra-canals” or “supernumerary canals”. In contrast, branches extending from an excretory canal were not considered to be an excretory canal, as they are secondary extensions not growing directly from the soma. The outgrowth and guidance defects of the excretory canals of HSPGs or other mutants have been studied elsewhere^4,21–23^. Here we focus on studying the uncommon supernumerary excretory canals defect, as it offers the unique opportunity to investigate the mechanisms that restrict the number of cellular extensions. The excretory cell is an ideal model to address this question as its development has been characterised in detail^11,24,25^, and its outgrowth and guidance are regulated by pathways shared with neurons^4,23,26^.

**Figure 1.**
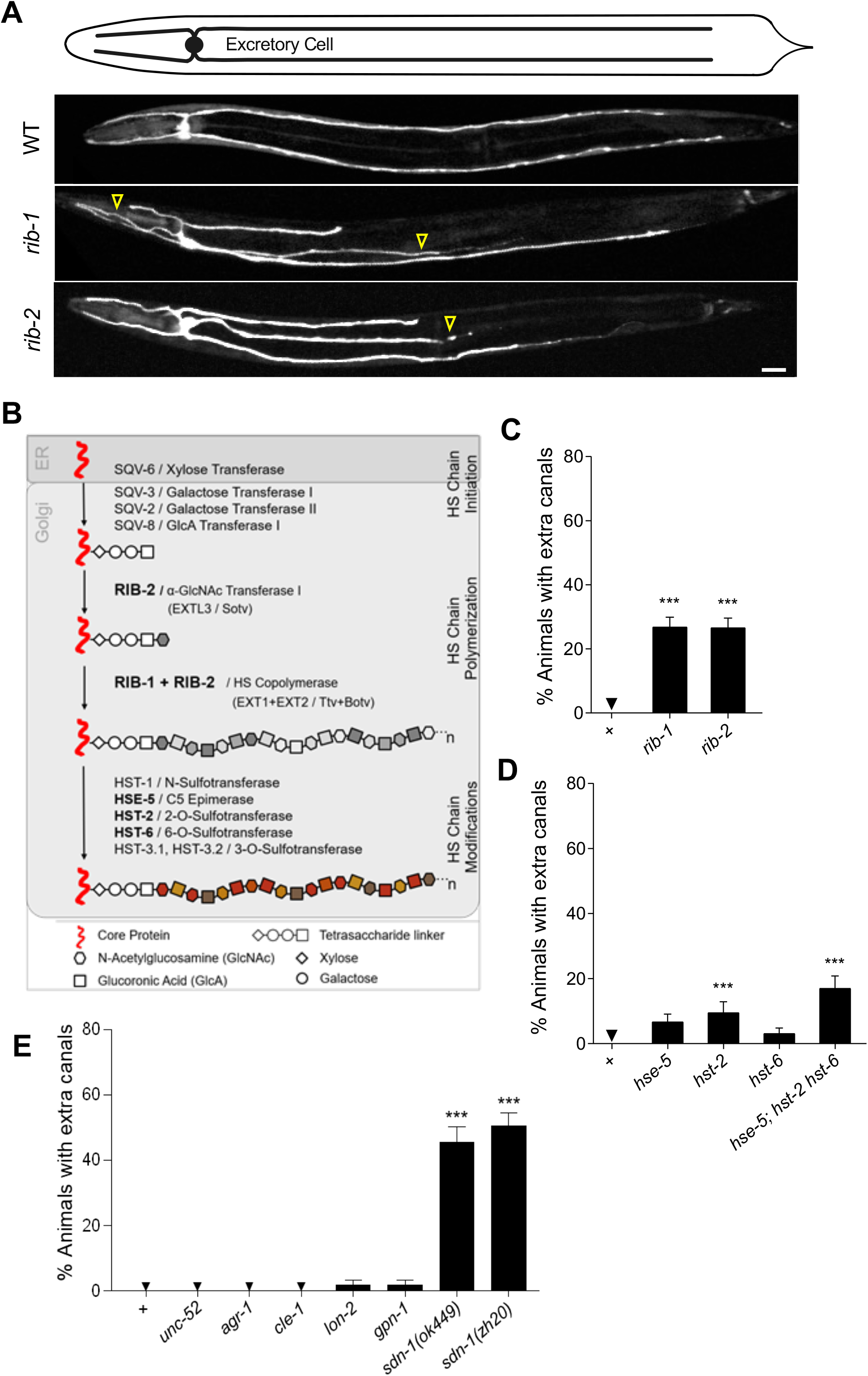
Perturbation of HSPGs results in additional cellular projections. (**A**) Ventral views of fourth larval stage animals by fluorescence microscopy. The excretory canal cell’s soma is located ventrally in the head. In the wild type, the excretory cell possesses 4 canals along the animal’s body sides, 2 anteriorly and 2 posteriorly. In mutants for the genes *rib*-*1* and *rib*-*2*, which encode the HS co-polymerase, the excretory cell displays supernumerary canals (i.e. more than 4 canals). Yellow arrowheads point supernumerary canals. Scale bar, 20 μm. (**B**) Schematics of HS chains biosynthesis on HSPG core proteins. Following the addition of a tetrasaccharide linker on specific Ser residues on the core proteins, HS polysaccharide chains are extended by the action of the HS copolymerase, which is composed of RIB-1 and RIB-2 in *C*. *elegans*. HS chains are subsequently modified by several modifying enzymes. (**C**) Quantification of the supernumerary canals of the excretory canal cell defects in hypomorphic mutants for the genes *rib*-*1* and *rib*-*2*. (**D**) Quantification of supernumerary excretory canals defects shown by mutants for genes encoding enzymes responsible of HS chain modifications. (**E**) Quantification of defective canal number in mutants for genes encoding HSPG core proteins. Animals mutant for *sdn*-*1/*syndecan, but not for other HSPGs, have significantly more than four canals. Error bars are standard error of the proportion. Asterisks denote significant difference: * p ≤ 0.05, *** p ≤ 0.001 (z-tests, p-values were corrected by multiplying by the number of comparisons). ns, not significant.

Having observed defects in the number of cellular projections in HS chain biosynthesis mutants, we next examined whether HS chain modification may be important in this process. Indeed, subsequent to their polymerization, HS chains are chemically modified, which impacts their role in neuronal migration and guidance^15,27–29^. We tested whether HS chain modifications carried out by the *C*. *elegans* sulfotransferases HST-2/HS2ST1 and HST-6/HS6ST1 and the epimerase HSE-5/GLCE may affect the number of excretory canals, by examining single null mutants for each of these enzymes, as well as triple mutant animals using the reporter P*glt*-*3::venus*. We found that animals mutant for *hse*-*5, hst*-*2* or *hst*-*6* develop extra canals (**Fig. 1D**), indicating that the HS chains modifications may play a role here. We constructed a triple mutant strain and detected no enhancement of the defect in animals mutant for all three genes *hse*-*5, hst*-*2* or *hst*-*6*, suggesting that all these HS modifications affect the development of the excretory cell in a similar way (**Fig. 1D**). Taken together these results suggest that HS chain biosynthesis and modification are required for the establishment of the proper number of canals in the excretory canal cell.

Disruption of the HS co-polymerase genes *rib*-*1* and *rib*-*2*, or of the HS modifying enzymes, affects the biosynthesis of all HSPGs simultaneously. To determine which HSPG(s) specifically act(s) to regulate the number of excretory canals, we systematically examined null or strong loss-of-function mutations that disrupt each of the HSPGs individually. Six HSPGs are known in *C*. *elegans*, namely UNC-52/Perlecan, AGR-1/Agrin, CLE-1/Collagen type XVIII, LON-2/Glypican, GPN-1/Glypican, and SDN-1/Syndecan. We found that among the HSPG core-protein mutants tested (**Fig. 1E**), only *sdn*-*1(zh20)* null mutants showed a significant number of extra canals (46% of the mutants; **Fig. 1E**) compared to wild type. Like for *rib*-*1* and *rib*-*2* mutants, the number and position of the extra canals vary in *sdn*-*1* mutants. A similarly penetrant defect was observed in animals mutant for a second allele of *sdn*-*1*, the strong loss-of-function *sdn-1(ok449)* (46%, **Fig. 1E**). Thus, the HSPG SDN-1/Syndecan limits the number of excretory canal cell projections.

### Wild-type and supernumerary canals display similar temporal and sub-cellular characteristics

At ∼ 400 min of embryonic development, the soma of the excretory canal cell undergoes lumen formation and two lateral extensions start projecting^24^ (**Fig. 2A**). At the 2-fold stage, these lateral extensions bifurcate to give rise to the four lateral canals, generating the cell’s typical H shape. These canals lengthen and their lumen extends along the length of the canals, until they span the entire length of the animal^24,26,30^. To determine when the supernumerary canals form in *sdn*-*1* mutants, we examined wild-type and *sdn*-*1(zh20)* mutant embryos at the time of canal formation (3-fold stage ≈ 550min of development). No extra canals are ever seen in wild-type embryos at this time or later; on the other hand, supernumerary canals are already present in *sdn*-*1* mutant embryos (**Fig. 2B**). Like normal canals, the supernumerary canals extend from the soma of the excretory canal cell, at the same time as normal canals, indicating that the mechanism at play acts during the initial development of the cell to regulate the number of cellular projections stemming from the soma. Furthermore, the penetrance of the defects in *sdn*-*1(zh20)* mutant animals is stable between the first (L1) and fourth (L4) larval stages (**Fig. 2C**), consistent with the notion that *sdn*-*1*/Syndecan functions during the initial development of the excretory canal cell to control canal number.

**Figure 2.**
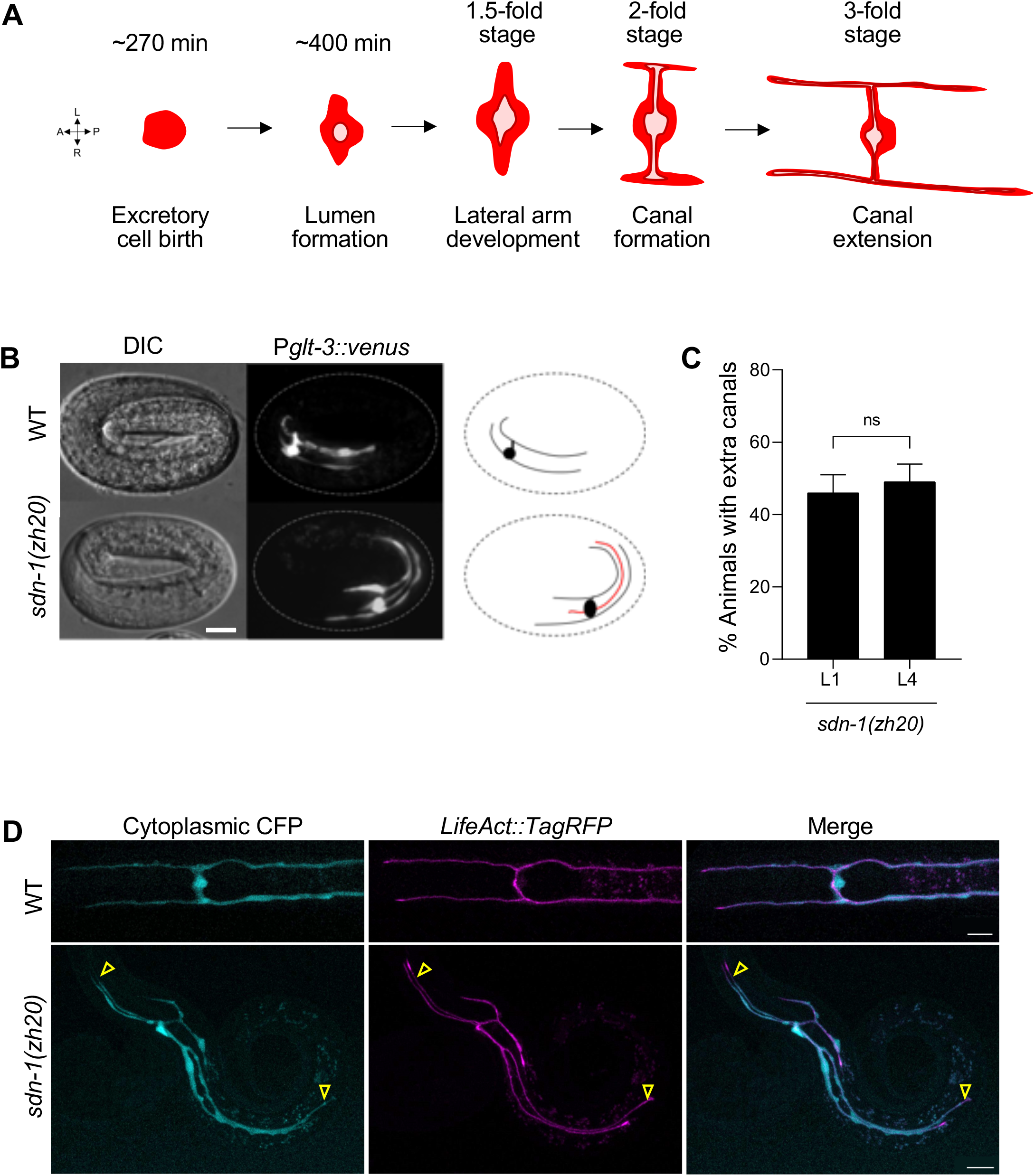
Supernumerary canals of the excretory canal cell displayed by *sdn*-*1*/Syndecan mutants develop embryonically. (**A**) Timeline of the excretory canal cell development. During embryonic development, the excretory cell precursor is born at ∼ 270 min, followed by tube formation starting at ∼ 400 min, and the formation of two lateral arms at the 1.5-fold stage, which will bifurcate to form four canals at the 2-fold stage. These excretory canals keep extending along the animal’s body side until the 4^th^ larval stage. (**B**) Fluorescence microscopy images of wild-type and *sdn*-*1(zh20)* mutant embryos. The excretory cell is visualized with marker P*glt*-*3::venus*. Supernumerary canals are already visible when the normal set of canals develops during embryogenesis. Panels to the right schematize the excretory canal cell with a presumptive supernumerary canals in red as we cannot differentiate it from the normal canals. (**C**) The supernumerary canal phenotype of *sdn*-*1(zh20)* null mutant animals is similar in penetrance at the 1^st^ (L1) and 4^th^ (L4) larval stages. (**D**) Confocal microscopy images of 1^st^ larval stage wild-type and *sdn*-*1(zh20)* animals. The cytoplasm of the excretory cell is visualized by the marker P*glt-3::cfp* (cyan) and actin by the marker *LifeAct::TagRFP* (magenta). All the excretory canals, including the supernumerary ones in *sdn*-*1* mutants, display actin apically in a similar manner. Arrowheads point to the tip of each supernumerary canals. Scale bar, 10 μm. Error bars are standard error of the proportion. ns, not significant (z-test).

To address whether the additional cellular projections are bona fide canals, we examined actin organization in the excretory cell using a *life*-*Act* transgene fused to the marker tagRFP (expressed specifically in the excretory canal cell). In wild-type animals, actin is present at the terminal web lining under the apical membrane along the lumen of the canals, surrounded by a cytoplasmic marker expressed in the excretory canal cell^23^ (P*glt*-*3*::CFP; **Fig. 2D**). Actin was similarly organized in the supernumerary canals of the *sdn*-*1(zh20)* mutant animals (**Fig. 2D**). Thus, the supernumerary canals observed in *sdn*-*1*/Syndecan mutants display similar temporal and sub-cellular characteristics to wild-type canals.

### Syndecan functions cell-autonomously to restrict the number of cell projections in the excretory canal cell

In a ∼ 500 min old wild-type embryo (Wormatlas^31,32^, **Fig. 3A**), when the excretory canals start projecting, the excretory cell is in direct contact with the pharynx and neurons, and in close proximity with the body wall muscles. Later on, the canals line up close to lateral epidermal seam cells. To address where SDN-1/Syndecan functions to regulate canal number, we tested where its expression may suffice for normal excretory canal cell development. For this, we generated transgenic *sdn*-*1(zh20)* animals that express wild-type copies of *sdn*-*1(+)* under several cell-specific promoters and analysed their capacity to rescue the extra canal defects of *sdn*-*1(zh20)* mutants. Expression in cells near the excretory cell such as the seam cells, body wall muscles, pharyngeal muscles, 6 mechanosensory neurons, or pan-neuronal expression did not rescue the defects of the *sdn*-*1(zh20)* mutants (**Fig. 3B**). Also, the two CAN neurons, which are in close proximity with the excretory canals all along the body sides and are physiologically related^33^, do not impact canal development since *unc*-*39(e257)* mutants, which are strongly defective in CAN cell migration and axon pathfinding^34^, have normal excretory canals (0%, N=119). In contrast, expression of wild-type copies of *sdn-1(+)* specifically in the excretory cell (transgene P*glt-3::sdn-1(+)*) strongly rescued the defects of the *sdn*-*1(zh20)* mutants (**Fig. 3B**), indicating that *sdn*-*1* functions in the excretory cell. The gene *sdn*-*1* is expressed in the excretory cell during its development^35^ (**Fig. S1A**), consistent with SDN-1/Syndecan functioning cell-autonomously to restrict the number of cell projections in the excretory canal cell.

**Figure 3.**
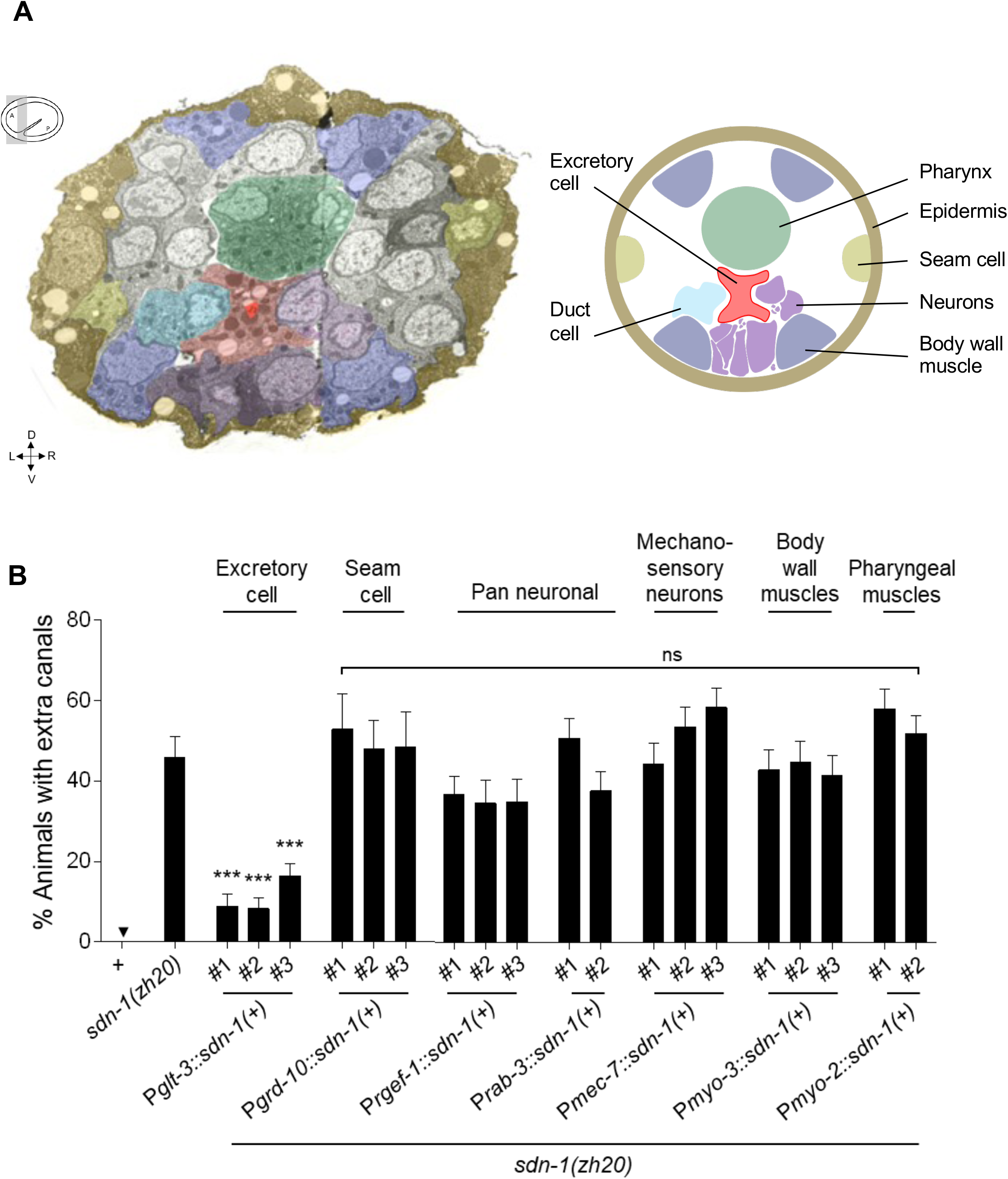
*sdn*-*1*/Syndecan acts in a cell-autonomous manner to control canal number during development. (**A**) Left panel is a transverse view of an approximately 500 min embryo, by transmission electron microscopy (TEM^1^). Cells surrounding the excretory cell (red) have been artificially coloured (epidermis in brown, seam cells in beige, body wall muscles in blue, neurons in purple, and excretory duct cell in cyan). Right panel is a schematic representation of the TEM view. (**B**) The supernumerary canal defects displayed by *sdn*-*1(zh20)* null mutant animals are fully rescued by expressing wild-type copies of *sdn*-*1*/syndecan under a heterologous promoter that drives expression specifically in the excretory cell [P*glt-3::sdn-1(+)*] indicating that *sdn*-*1* functions cell-autonomously to control the number of canals. Expression of wild-type copies of *sdn*-*1*/syndecan under other heterologous promoters driving expression in neighbouring tissues did not rescue the defects of *sdn*-*1(zh20)* mutants. Error bars are standard error of the proportion. Asterisks denote significant difference compared to *sdn*-*1(zh20)* group: *** p ≤ 0.001 (z-tests, p-values were corrected by multiplying by the number of comparisons). ns, not significant.

### Syndecan and Netrin function in same pathway to restrict the number of cell projections in the excretory canal cell

To gain insight into the mechanism controlling cell projection number, we determined whether pathways known to guide the migration of developing canals^4^ might also function to regulate canal number. Hedgcock and colleagues anecdotally reported supernumerary canals^4^ in mutants for *unc*-*6*/Netrin and its receptors. UNC-6/Netrin signals can be attractive in cell migration and/or axon pathfinding (**Fig. 4A**), in which case UNC-6 interacts with two UNC-40 receptors, inducing cytoskeleton rearrangements to orient migration towards the UNC-6 signal^4–6^. UNC-6/netrin signals can also be repulsive, where UNC-6 links a heterodimer of an UNC-40 receptor and an UNC-5 receptor, leading to migration away from the UNC-6/netrin signal^7^(**Fig. 4A**). We thus tested whether the guidance cue UNC-6/Netrin and its receptors UNC-40/DCC and UNC-5/UNC5, are involved in the regulation of excretory canal number. Loss of function of *unc*-*6*, with either the null mutation *unc-6(ev400)* or the hypomorphic mutation *unc-6(e78)*, leads to supernumerary canals on the excretory cell (**Fig. 4B and C**). Furthermore, loss of function of either of the *unc*-*6*/netrin receptors, using single null mutations *unc*-*40(e1430)*, *unc*-*40(e271)* or *unc*-*5(e53)*, also results in supernumerary canals (**Fig. 4B and C**), and the simultaneous loss of both receptors in *unc-40(e1430) unc-5(e53)* double mutants does not enhance the defects, pointing to their action in the same mechanism to control cell projection number in the excretory canal cell. Taken together, these results indicate that the netrin guidance cue as well as its main receptors UNC-40/DCC and UNC-5/UNC5, are involved in a mechanism controlling the number of cellular extensions developed.

**Figure 4.**
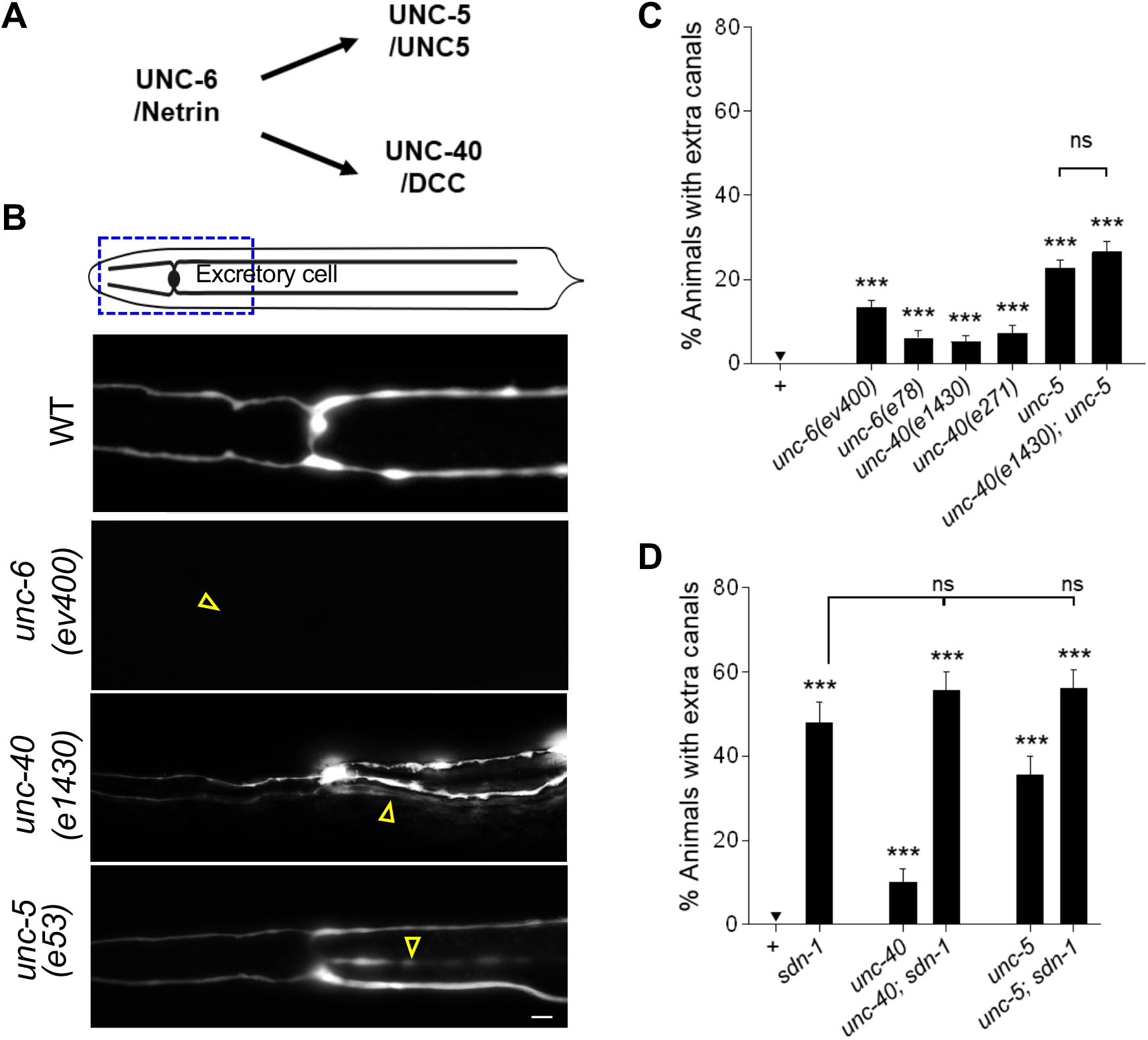
Disruption of the UNC-6/Netrin guidance pathway affects the excretory cell canal number. (**A**) The UNC-6/Netrin signal acts through its main receptors UNC-40/DCC and UNC-5/UNC5. (**B**) Ventral views of 4^th^ larval stage wild-type and mutant animals, where the excretory cell is visualized by expressing *Pglt*-*3::venu*s. The region observed in these images is indicated by the rectangle on the diagram. Arrowheads point supernumerary canals. Scale bar, 10 μm. (**C**) Quantification of supernumerary canals defects in mutants for genes encoding guidance cue molecules or their receptors. Null mutant animals *unc-6(ev400)*, *unc*-*40(e1430)* and *unc*-*5(e53*) show supernumerary canals. (**D**) Quantification of supernumerary canals defects in animals doubly mutants for syndecan/*sdn*-*1(zh20)* and genes encoding guidance receptors, *unc*-*40(e1430)* or *unc*-*5(e53)*. The fact that these double mutants are not more severe than the single mutant *sdn*-*1(zh20)* suggests that the genes function in the same pathway. Error bars are standard error of the proportion. Asterisks denote significant difference: *** p ≤ 0.001, ** p ≤ 0.01 and * p ≤ 0.05 (z-tests, p-values were corrected by multiplying by the number of comparisons). ns, not significant.

To ask whether the *unc*-*6*/*netrin* receptors contribute to the same mechanism as *sdn*-*1*/Syndecan in controlling cell projection number of the excretory canal cell, we analysed their genetic interactions using null alleles. Simultaneous loss of *sdn*-*1* and either *unc-5* or *unc*-*40*, in the double mutants *sdn-1(zh20); unc-5(e53)* or *sdn*-*1(zh20); unc*-*40(e1430)* did not enhance the supernumerary canals phenotype (**Fig. 4D**). This is consistent with the notion that *sdn*-*1*/Syndecan and the two guidance receptors function in the same pathway and suggests that they may participate in a common mechanism with SDN-1/Syndecan to regulate the number of cell projections.

### MIG-2/RhoG and CED-10/Rac are required for the formation of extra projections upon SDN-1/Syndecan loss

The Rho-GTPases MIG-2/RhoG and CED-10/Rac are molecular switches that control cytoskeleton dynamics during embryonic morphogenesis^11^ and axonal pathfinding^10^, playing roles in the UNC-6/Netrin pathway and cooperating with HSPGs like Syndecan in mammal cells^36–38^. To investigate the role of these Rho-GTPases in the control of cell projection number, we used mutations in these genes. Single mutant animals with the partial loss-of function mutation *ced*-*10(n1993)* or the null mutation *mig*-*2(mu28)* show no supernumerary canals (**Fig. 5A,B**). The double mutants *sdn*-*1(zh20)*; *ced*-*10(n1993)* and *mig-2(mu28) sdn-1(zh20)*, however, both display a reduced number of supernumerary canals compared to the single mutant animals *sdn*-*1(zh20)*. This suggests that the normal functions of CED-10/Rac and of MIG-2/RhoG promote canal projection, and are required for the formation of extra projections upon SDN-1/Syndecan loss.

**Figure 5.**
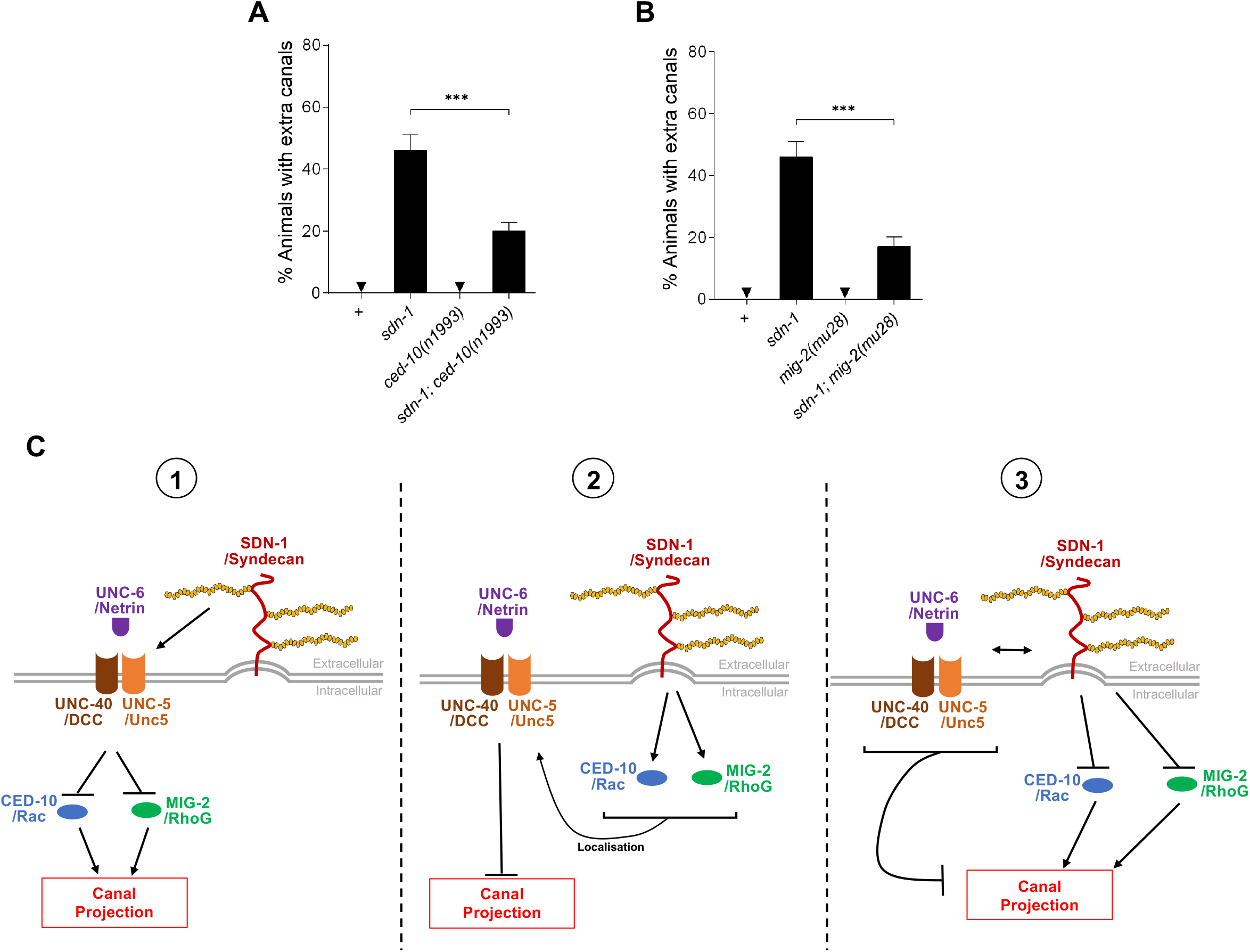
The control of the number of canals requires Rho GTPases. (**A**) Quantification of supernumerary canal defects in mutants of *ced*-*10*. Alone, the *ced*-*10(n1993)* loss-of-function mutants do not induce extra canals, but suppresses the canal number defects *sdn*-*1(zh20)* null mutants. (**B**) Quantification of supernumerary canal defects in mutants of *mig-2*. Single null mutants *mig*-*2(mu28)* alone does not display supernumerary canals, but suppresses the canal number defects *sdn*-*1(zh20)* null mutants. (**C**) Schematic summary of molecules implicated in controlling cellular projection number and possible mechanisms. SDN-1/Syndecan is central HSPG to control the number of canals in the excretory cell and could coordinate signals from the UNC-6/Netrin guidance pathway to drive proper cell development. The mechanism under the scope may converge onto RhoGTPases to regulate canal number. Error bars are standard error of the proportion. Asterisks denote significant difference: *** p ≤ 0.001 (z-tests, p-values were corrected by multiplying by the number of comparisons). ns, not significant.

## DISCUSSION

Cells with complex morphologies such as neurons require establishment of precise cell shapes, with a defined number of cellular projections, to enable function. Understanding of outgrowth and guidance of cellular projections has greatly progressed, and HSPGs have emerged as important players in these morphogenetic events^13,15,16,39^. In contrast, the regulation of the number of cellular projections at the very onset of their development is poorly understood. Thanks to a new phenotype where a developing cell has more projections than it should, we were able to identify a mechanism by which the number of extensions established during development is regulated. We show that early in the development of the excretory canal cell HSPG SDN-1/Syndecan acts in a cell-autonomous manner, together with the netrin guidance pathway and two Rho-GTPases CED-10/Rac and MIG-2/RhoG to control the number of excretory canals (**Fig. 5C**).

### Study of a novel phenotype: A polarized cell developing supernumerary extensions

Several defects of the excretory canal cell observed in HSPG mutants were previously studied (shorter or missing canals likely due to outgrowth issues^4,11,21,22^, guidance defects^4,16^ or branched canals^22^). Here, we report a new phenotype, which had been anecdotally noted by others previously^4,40^, but remained unstudied. The depletion of HS biosynthesis in *rib*-*1* and *rib*-*2* mutants, the loss of *sdn*-*1*/Syndecan, and the disruption of the *unc*-*6*/Netrin pathway, all induce supernumerary cellular projections. These supernumerary projections stem directly out of the soma of the excretory canal cell (**Fig. 1A**) and display temporal and cellular hallmarks of wild-type canals (**Fig. 2B**). Given that no extra canals were ever seen at that time in wild-type embryos, these supernumerary extensions do not appear to arise from uneliminated transitional protrusions that would later be eliminated in wild-type development, and instead appear to arise as *de novo* projections when the mechanism to restrict them is abrogated. This is in contrast with the extension of short branches from a pre-existing cellular projection^16,41^, such as in aging neurons^42–44^, or the bifurcation of a growth cone leading to multiple extensions^45^. The growth of multiple cellular projections having the hallmarks of mature extensions and growing from a cellular soma has been observed only rarely in cells in culture under specific conditions^12^ and it even more rarely *in vivo*^46^. Thus, the model described here sets the ground for detailed studies of the molecular and cellular mechanisms at play.

### Syndecan restricts the number of canals in the excretory cell

Our systematic analysis of all available HSPG core protein mutants in *C*. *elegans* led to the identification of one key evolutionarily conserved HSPG, SDN-1/Syndecan, as a cell-autonomously acting inhibitor of supernumerary projections (**Fig. 1E**, **Fig. 3B**). Whereas shedding of extracellular regions of Syndecan has been reported in other contexts^47^ allowing it to act at a distance, in the mechanism of cellular projection control in the excretory canal cell, Syndecan was found to function in this very cell during development, as only expression in the developing excretory cell restored function, while expression in neighbouring cells did not (**Fig. 3B**).

HSPGs function can be mediated by either their core protein or their glycosaminoglycan HS chains^15^, particularly during morphogenesis of polarized cells^16^. Mutation of the HS chain polymerisation in *rib*-*1* and *rib*-*2* mutants alters the number of cellular projections (**Fig. 1C**), indicating that either the HS chains per se play a role in this mechanism, or that the level or targeting of the entire HSPG may be affected. Similar penetrance of the supernumerary canal (**Fig. 1E**) was seen in the null mutant *sdn*-*1(zh20)* and the partial mutant *sdn*-*1(ok499)*, which expresses a truncated SDN-1/Syndecan protein lacking the HS chains attachment region^48^. This supports the notion that the HS chains are important for the role of Syndecan in the control of cell projection number, either because they play a direct role in the mechanism as for axonal guidance^16,49,50^, or because the absence of HS chains impacts the level or targeting of SDN-1. Similarly, disruption of HS chain maturation in the triple HS modifying enzymes null mutant *hse*-*5(tm472); hst*-*2(ok595) hst*-*6(ok273)* showed supernumerary cellular projections (**Fig. 1D**), suggesting that HS chain maturation is important for HSPGs in the regulation of cellular projection number.

### Syndecan cooperates with guidance pathways to control the number of cellular projections

We showed that the guidance cue UNC-6/Netrin and its receptors UNC-40/DCC and UNC-5/UNC5 are part of the mechanism restricting the number of cellular projections of the excretory cell (**Fig. 4C**). HSPGs have been described as co-receptors mediating interactions of guidance cues with their receptors^13,15^, and here we uncover that SDN-1/Syndecan and UNC-6/Netrin receptors UNC-40/DCC and UNC-5/UNC5 function in same pathway to control cellular projection number in the developing excretory canal cell. Consistent with our findings, a link between Syndecan and the UNC-6/Netrin pathway was suggested in the context of D-type motor axon guidance^14^, distal tip cell migration^51^, and in synaptic development, where the content of UNC-40/DCC at chemical synapses requires SDN-1/Syndecan^52^, reinforcing the notion that Syndecan and the Netrin guidance pathway together underlie key developmental events.

### Syndecan controls the number of cell extension through two Rho-GTPases

Similarities in the cytoskeletal organisation exist between the growth cone of a developing neuron^11^ and the tip of a growing canal, where actin and microtubules need to be tightly regulated for proper development^11,25^. CED-10/Rac and MIG-2/RhoG are two Rho-GTPases that regulate cytoskeletal organisation in numerous biological processes^10,11^. Of particular interest, they regulate the actin cytoskeleton downstream of the UNC-6/Netrin guidance pathways in the growth cone of axons^6,53^. We have showed here that MIG-2 and CED-10 are required for the development of supernumerary cellular projections when the SDN-1-mediated mechanisms to control the number of excretory canals is disrupted, establishing a novel link between these Rho-GTPases and a HSPG in *C*. *elegans*. Several scenarios for the role of SDN-1/Syndecan can be envisaged. (1) SDN-1/Syndecan may act as a co-receptor with the UNC-6/Netrin receptor(s), modulating the function of downstream targets of this pathway such as the cytoskeleton regulators CED-10/Rac and MIG-2/RhoG, thus controlling the number of extensions formed by the cytoskeleton regulation. (2) SDN-1/Syndecan may directly act on the Rho-GTPases CED-10/Rac and MIG-2/RhoG, as mammalian Syndecans do^36–38^, to modulate the function of the UNC-6/Netrin pathway by regulating UNC-40/DCC plasma membrane localization, as observed in axon development^54^. (3) SDN-1/Syndecan might act as a central regulator, impacting the UNC-6/Netrin pathway as well as the cytoskeleton modulators CED-10/Rac and MIG-2/RhoG in distinct ways that converge in the regulation of cell projection number during the development of the excretory canal cell.

Taken together our results uncover a novel developmental mechanism ensuring the control of cellular projection number during the development of highly polarized cells. HSPG SDN-1/Syndecan is central to this mechanism, which also employs the guidance cue UNC-6/Netrin and its main receptors UNC-40/DCC and UNC-5/UNC5 in the same genetic pathway, as well as two Rho-GTPases CED-10/Rac and MIG-2/RhoG. Further studies are needed to address the impact of SDN-1/Syndecan on the regulation of receptor localization and of cytoskeleton dynamics during cell development. Given the evolutionary conservation of developmental mechanisms and the molecules implicated, this work provides information relevant to understanding the cellular and molecular bases of the development of precise cellular morphologies in varied cell types across animals.

## ACKNOWLEGMENTS

We thank Marianne Moore for her technical assistance at the early stages of the project; Denis Flipo (UQAM) for his confocal microscopy expertise; several investigators for their gift of plasmids, including Dr. A. Fire (pPD clones), Dr. Y. Kohara (yk cDNA clones); Wormbase; the *Caenorhabditis* Genetics Center, which is funded by NIH Office of Research Infrastructure Programs (P40 OD010440) for strains; WormAtlas and WormImage, which are funded by NIH OD010943 to David H. Hall; John White and Jonathan Hodgkin for donation of the MRC/LMB archives to the D. Hall laboratory for curation; as well as Richard Durbin, who did the original TEM work on the embryo, which the Hall laboratory published for him on WormAtlas. This work was supported by the National Science and Engineering Research Council of Canada, the Fond de Recherche du Québec-Santé (C.B. Research Scholar Jr2 and Sr), the Canadian Funds for Innovation-John Evan Leaders Equipment Grant, two Excellence in Doctoral Research Scholarships from the Research Center CERMO-FC (Centre d’excellence de recherche sur les maladies orphelines-Fondation Courtois) to R.D., three Excellence Masters Research Scholarships to M.B.T by the UQAM Fondation.

