## Supplementary Methods and Files for "Syndecan Regulates Cellular Morphogenesis in Cooperation with the Netrin Guidance Pathway and Rho-family GTPases"

**S1 Table.** List of primers used.

| Gene | Primer | Sequence | PCR product (bp) | Cosmid Coordinates |
|---|---|---|---|---|
| ***rib-1(qm32)* sequencing** | | | | |
| | oCB1026 | gggtgcgtaaggagatgagg | 456 | F12F6 39341…39360, forward |
| | oCB1027 | ggcaaccagccatcacagcc | | F12F6 39797…39816, reverse |
| ***rib-2(qm46)* sequencing** | | | | |
| | oCB1028 | caacttatcggatcttcaacc | 437 | K01G5 4053…4073, forward |
| | oCB1029 | ttccagcggtccaaggagg | | K01G5 4508...4490, reverse |
| ***hse-5(tm472)*** | | | | |
| Mutant specific | oCB1055 | atcgtgtacgatgtgtcagc | 546 | B0285 15457…15476, forward |
| | oCB1056 | attcgcctcatacggtttcc | | B0285 16217…16236, reverse |
| Wild-type specific | oCB1055 | atcgtgtacgatgtgtcagc | 779 | B0285 15457…15476, forward |
| | oCB1057 | aactttctctcggcaattg | | B0285 17233…17251, reverse |
| ***hst-2(ok595)*** | | | | |
| Mutant specific | oCB1052 | tattacaacatggacggagc | 692 | C34F6 17093…17112, forward |
| | oCB1054 | aacattatgcgcatgaacgc | | C34F6 15085…15104, reverse |
| Wild-type specific | oCB1052 | tattacaacatggacggagc | 486 | C34F6 17093…17112, forward |
| | oCB1053 | ttagcagtgattcaattacg | | C34F6 16626…16645, reverse |
| ***hst-6(ok273)*** | | | | |
| Mutant specific | oCB1049 | ttagacgtggctgttctcac | 723 | Y34B4A 39244…39263, forward |
| | oCB1051 | tgtgagtctgttaagggtgg | | Y34B4A 41012…41031, reverse |
| Wild-type specific | oCB1049 | ttagacgtggctgttctcac | 804 | Y34B4A 39244…39263, forward |
| | oCB1212 | agaaatgttgtgtagaagtag | | Y34B4A 40028…40048, reverse |
| ***sdn-1(zh20)*** | | | | |
| Mutant specific | oCB837 | aaagagatgccggtcaggtg | 410 | F57C7 28510…28529, forward |
| | oCB842 | aatggacgggatgagtgtcc | | F57C7 26861…26880, reverse |
| Wild-type specific | oCB837 | aaagagatgccggtcaggtg | 293 | F57C7 28510…28529, forward |
| | oCB876 | cttcagattcgagcctgctttgc | | F57C7 28237…28259, reverse |
| ***sdn-1(ok449)*** | | | | |
| Mutant specific | oCB1114 | ttctgcctgtcgacttactc | 297 | F57C7 28453…28472, forward |
| | oCB1125 | aagattgcggtaaaacacatc | | F57C7 27692…27712, reverse |
| Wild-type specific | oCB1114 | ttctgcctgtcgacttactc | 478 | F57C7 28453…28472, forward |
| | oCB1115 | ttcgtcgtcggttgggtagc | | F57C7 27994…28013, reverse |
| ***gpn-1(tm595)*** | | | | |
| Mutant specific | oCB1309 | agtcgattgcaaacgaatacg | 468 | F59D12 36353…36373, forward |
| | oCB1310 | tcacacagtacgcttggcacg | | C03H12 356…376, reverse |
| Wild-type specific | oCB1309 | agtcgattgcaaacgaatacg | 393 | F59D12 36353…36373, forward |
| | oCB1311 | aagctttccatgcatactcgc | | C03H12 1822…1842, reverse |
| ***agr-1(tm2051)*** | | | | |
| Mutant specific | oCB891 | cgaaaaatcgagagcaaaagg | 362 | F41G3 22649…22669, forward |
| | oCB893 | tcagattcttgacacatccc | | F41G3 21037…21056, reverse |
| Wild-type specific | oCB890 | tttgaactcttggacgaacc | 1210 | F41G3 22308…22327, reverse |
| | oCB891 | cgaaaaatcgagagcaaaagg | | F41G3 22649…22669, forward |
| ***cle-1(cg120)*** | | | | |
| Mutant specific | oCBQc45 | ggtactggacatggatctgg | 751 | F39H11 14593…14612, reverse |
| | oCBQc46 | tgccaaactcgcttatctgg | | F39H11 11859…11878, forward |

| | | | | |
|---|---|---|---|---|
| Wild-type specific | oCBQc47 | gctttcgagtaatgtacagg | 879 | F39H11 12718…12737, reverse |
| | oCBQc46 | tgccaaactcgcttatctgg | | F39H11 11859…11878, forward |
| ***unc-6(ev400)*** | **ARMS-PCR** | | | |
| Mutant specific | oCB1712 | gacacttgtgatgctagaaaccatt**g**c**t** | 217 | F41C6 21346…21374, Forward inner |
| | oCB1711 | cagacttgtaaagtgccattgaatctgg | | F41C6 21536…21563, Reverse outer |
| Wild-type specific | oCB1713 | ttgaaaaggttgtcgcccaaaagta**t**ac | 198 | F41C6 21202…21229, Forward outer |
| | oCB1714 | gttagaagagaggctggatgggagtg | | F41C6 21374…21399, Reverse inner |
| Outer primers | oCB1711 + oCB1713 | | 362 | |
| ***unc-40(e1430)*** | | | | |
| For sequencing | oCB1078 | agagaccagggagttacagg | 367 | T19B4 12759…12778, forward |
| | oCB1079 | atcaatgcgctgtacatgtg | | T19B4 13106…13125, reverse |
| ***unc-5(e53)*** | **ARMS-PCR** | | | |
| Mutant specific | oCB1757 | gatgacaactcgaagagcaagcac**g**c**t** | 280 | B0273 15164…15190, forward inner |
| | oCB1760 | aatttcagttgacggtggttggtccgaatg | | B0273 15415…15444, reverse outer |
| Wild-type specific | oCB1758 | gatggaggatggagttcatggagtga**g**tg | 126 | B0273 15190…15218, reverse inner |
| | oCB1759 | tcagatcatctccaaaacatggctgtcc | | B0273 15092…152119, forward outer |
| Outer primers | oCB1759 + oCB1760 | | 353 | |
| ***ced-10(n1993)*** | **SS-PCR** | | | |
| Mutant specific | oCB2019 | ccgccacaaagagccaaa**atctcgttca** gtacgg**g** | 369 | C09G12 34086…34120, reverse mutant specific |
| | oCB2017 | tagagagtggaaacgcccag | | C09G12 33752…33771, forward |
| Wild-type specific | oCB2018 | ccgccacaaagagccaaa**atctcgttca** gtacggt | 369 | C09G12 34086…34120, reverse wild-type specific |
| | oCB2017 | tagagagtggaaacgcccag | | C09G12 33752…33771, forward |
| ***mig-2(mu28)*** | **ARMS-PCR** | | | |
| Mutant specific | oCB1717 | ggaacgttgtgaacttaggattg**g**g**a** | 539 | C35C5 20375…20400, forward inner |
| | oCB2088 | cacattgaacctctgcactttacg | | C35C5 20619…20642, reverse outer |
| Wild-type specific | oCB1718 | cataatcctcctgtccagcagta**g**cc | 268 | C35C5 20400…20425, reverse inner |
| | oCB2087 | aacgccctcttcttgagtatttcg | | C35C5 19887…19910, forward outer |
| Outer primers | oCB2087 + oCB2088 | | 756 | |

**Underlined and Bold** bases correspond to mismatch compared to the sequence allowing single nucleotide genotyping using ARMS-PCR[1] or SS-PCR[2].


1. Little, S. Amplification-refractory mutation system (ARMS) analysis of point mutations. *Curr Protoc Hum Genet* **Chapter 9**, Unit 9.8 (2001).
2. Touroutine, D. & Tanis, J. E. A Rapid, SuperSelective Method for Detection of Single Nucleotide Variants in Caenorhabditis elegans. *Genetics* **216**, 343–352 (2020).

**S2 Table**. List of strains used.

| Strain | Genotype | Transgene | Reference |
|---|---|---|---|
| N2 | | | (1) |
| VQ1157 | *arIs164* V | [P*glt-3::Venus*] | (2) |
| | *bgIs312* | *bgIs312* [P*pes-6::GFP*] | |
| **HS chain biosynthesis, HS and CS modifying enzymes** | | | |
| VQ342 | *rib-1(qm32)* IV*; bgIs312* | | (3) |
| VQ343 | *rib-2(qm46)* III*; bgIs312* | | (3) |
| VQ1322 | *hst-2(ok595)* X*; arIs164* V | | This study |
| VQ1373 | *hse-5(tm472)* III*; arIs164* V | | This study |
| VQ1374 | *hst-6(ok273)* X*; arIs164* V | | This study |
| VQ1356 | *hse-5(tm472)* III*; hst-2(ok595) hst-6(ok273)* X*; arIs164* V | | This study |
| **Heparan Sulfate Proteoglycans** | | | |
| VQ1183 | *sdn-1(zh20)* X*; arIs164* V | | This study |
| VQ1230 | *sdn-1(ok449) X; arIs164 V* | | This study |
| VQ1170 | *lon-2(e678)* X*; arIs164* V | | This study |
| VQ1171 | *gpn-1(tm595)* X*; arIs164* V | | This study |
| VQ1217 | *agr-1(tm2051)* II*; arIs164* V | | This study |
| VQ1166 | *unc-52(e444)* II*; arIs164* V | | This study |
| VQ1184 | *cle-1(cg120)* I*; arIs164* V | | This study |
| **Guidance molecules and receptors** | | | |
| VQ1280 | *unc-6(ev400)* X*; arIs164* V | | This study |
| VQ1431 | *unc-6(e78)* X*; arIs164* V | | This study |
| VQ1765 | *unc-40(e1430)* I*; arIs164* V | | This study |
| VQ1471 | *unc-40(e271)* I*; arIs164* V | | This study |
| VQ1806 | *unc-5(e53)* IV*; arIs164* V | | This study |
| VQ1818 | *unc-40(e1430)* I*; unc-5(e53)* IV*; arIs164* V | | This study |
| VQ1577 | *unc-40(e1430)* I*; sdn-1(zh20)* X*; arIs164* V | | This study |
| VQ1467 | *unc-5(e53)* IV*; sdn-1(zh20)* X*; arIs164* V | | This study |
| **Intracellular effectors** | | | |
| VQ1730 | *ced-10(n1993)* IV; *arIs164* V | | This study |
| VQ1740 | *ced-10(n1993)* IV; *sdn-1(zh20)* X; *arIs164* V | | This study |
| QN68 | *mig-2(mu28)* X*; arIs164* V | | This study |
| VQ1462 | *sdn-1(zh20) mig-2(mu28)* X*; arIs164* V | | This study |

1. Brenner S. The genetics of Caenorhabditis elegans. Genetics. 1974 May;77(1):71–94.

2. Shaye DD, Greenwald I. The Disease-Associated Formin INF2/EXC-6 Organizes Lumen and Cell Outgrowth during Tubulogenesis by Regulating F-Actin and Microtubule Cytoskeletons. Developmental Cell. 2015 Mar;32(6):743–55.

3. Blanchette CR, Thackeray A, Perrat PN, Hekimi S, Bénard CY. Functional Requirements for Heparan Sulfate Biosynthesis in Morphogenesis and Nervous System Development in C. elegans. PLoS Genet [Internet]. 2017 Jan 9 [cited 2019 Nov 5];13(1). Available from: https://www.ncbi.nlm.nih.gov/pmc/articles/PMC5221758/

**S3 Table**. List of mutant alleles used.

| Gene | Allele | Nature of allele | Reference |
|---|---|---|---|
| ***rib-1*** | *qm32* | Stop codon converted into a Lys codon, likely extending the open reading frame into the 3'UTR. Partial loss of function. | (1) |
| ***rib-2*** | *qm46* | Arg to Gln amino acid substitution at residue 434. Partial loss of function. | (1) |
| ***hse-5*** | *tm472* | 1249 bp deletion and addition of an adenosine, which deletes most of exons 4 – 7 and generates a frame shift after exon 4. Null. | (2) |
| ***hst-2*** | *ok595* | 1336 bp deletion, deleting from exon 4 to part of exon 7. Predicted null. | (2) |
| ***hst-6*** | *ok273* | 1064 bp deletion, with the addition of nucleotides CTTT, which deletes exons 4 and 5 and generates a frameshift after exon 3. Predicted null. | (2) |
| ***sdn-1*** | *zh20* | 1258 bp deletion. No transcripts detected. | (3) |
| | *ok449* | 483 bp deletion in frame, lacks two glycosylation sites. Partial loss of function. | (4) |
| ***lon-2*** | *e678* | ~9 kb deletion. Null. | (5) |
| ***gpn-1*** | *tm595* | 1411 bp deletion, deletes part of exon 2, exon 3, and introduces early Stop codons. Likely null. | (6,7) |
| ***agr-1*** | *tm2051* | 423 bp deletion, deletes exons 26 and 27 resulting in an in-frame loss of 42 amino acids. | (8) |
| ***unc-52*** | *e444* | Arg to early stop codon substitution in exon 18. Partial loss of function. | (9) |
| ***cle-1*** | *cg120* | ~2 kb deletion, removes exons 18-20 and add premature termination. Partial loss of function. | (10) |
| ***unc-6*** | *ev400* | Gln to early stop codon substitution at the residue 78. Null. | (11) |
| | *e78* | Cys to Tyr substitution at the residue 410. Partial loss of function. | (12) |
| ***unc-40*** | *e271* | Arg to early stop codon substitution at the residue 824. Null. | (13) |
| | *e1430* | Arg to early stop codon substitution at the residue 157. Likely null. | (14) |
| ***unc-5*** | *e53* | Trp to early stop codon substitution at the residue 283. Null. | (15) |
| ***ced-10*** | *n1993* | Val to Gly substitution at the residue 190. Prenylation site affecting membrane targeting, partial loss of function. | (16) |
| ***mig-2*** | *mu28* | Trp to early stop codon substitution at the residue 60. Predicted null. | (17) |

15. Killeen M, Tong J, Krizus A, Steven R, Scott I, Pawson T, et al. UNC-5 Function Requires Phosphorylation of Cytoplasmic Tyrosine 482, but Its UNC-40-Independent Functions also

Require a Region between the ZU-5 and Death Domains. Developmental Biology. 2002 Nov 15;251(2):348–66.

15. Killeen M, Tong J, Krizus A, Steven R, Scott I, Pawson T, et al. UNC-5 Function Requires Phosphorylation of Cytoplasmic Tyrosine 482, but Its UNC-40-Independent Functions also

Require a Region between the ZU-5 and Death Domains. Developmental Biology. 2002 Nov 15;251(2):348–66.

16. Reddien PW, Horvitz HR. CED-2/CrkII and CED-10/Rac control phagocytosis and cell migration in Caenorhabditis elegans. Nature Cell Biology. 2000 Mar;2(3):131–6.

17. Zipkin ID, Kindt RM, Kenyon CJ. Role of a New Rho Family Member in Cell Migration and Axon Guidance in C. elegans. Cell. 1997 Sep 5;90(5):883–94.

**S4 Table.** Canal number defects in mutants and transgenic lines for each figure.

| Genotype | Transgene | N | % Defective | s.e.p. |
|---|---|---|---|---|
| **Figure 1C** | | | | |
| *bgIs312* | | 211 | 0 | 0 |
| *rib-1(qm32); bgIs312* | | 198 | 27 | 3.1 |
| *rib-2(qm46); bgIs312* | | 218 | 26 | 3.0 |
| **Figure 1D** | | | | |
| *arIs164* | | 156 | 0 | 0 |
| *hse-5(tm472); arIs164* | | 105 | 7 | 2.4 |
| *hst-2(ok595); arIs164* | | 116 | 9 | 2.6 |
| *hst-6(ok273); arIs164* | | 98 | 3 | 1.7 |
| *hse-5(tm472); hst-2(ok595) hst-6(ok273); arIs164* | | 100 | 17 | 3.8 |
| **Figure 1E** | | | | |
| *arIs164* | | 120 | 0 | 0 |
| *unc-52(e444); arIs164* | | 46 | 0 | 0 |
| *agr-1(tm2051); arIs164* | | 91 | 0 | 0 |
| *cle-1(cg120); arIs164* | | 63 | 0 | 0 |
| *lon-2(e678); arIs164* | | 101 | 2 | 1.4 |
| *gpn-1(tm595); arIs164* | | 106 | 2 | 1.3 |
| *sdn-1(ok449); arIs164* | | 112 | 46 | 4.7 |
| *sdn-1(zh20); arIs164* | | 159 | 49 | 4.0 |
| **Figure 2C** | | | | |
| *sdn-1(zh20); arIs164* **L1** | | 94 | 46 | 4.9 |
| *sdn-1(zh20); arIs164* **L4** | | 101 | 41 | 5.1 |
| **Figure 3B** | | | | |
| *arIs164* | | 158 | 0 | 0 |
| *sdn-1(zh20); arIs164* | | 170 | 51 | 3.8 |
| *sdn-1(zh20); arIs164; qvEx437* | P*glt-3::sdn-1* | 101 | 9 | 2.8 |
| *sdn-1(zh20); arIs164; qvEx438* | P*glt-3::sdn-1* | 108 | 8 | 2.7 |
| *sdn-1(zh20); arIs164; qvEx509* | P*glt-3::sdn-1* | 152 | 16 | 3.0 |
| *sdn-1(zh20); arIs164; qvEx352* | P*grd-10::sdn-1* | 34 | 53 | 8.6 |
| *sdn-1(zh20); arIs164; qvEx353* | P*grd-10::sdn-1* | 50 | 48 | 7.1 |
| *sdn-1(zh20); arIs164; qvEx354* | P*grd-10::sdn-1* | 33 | 48 | 8.7 |
| *sdn-1(zh20); arIs164; qvEx511* | P*rgef-1::sdn-1* | 114 | 37 | 4.5 |
| *sdn-1(zh20); arIs164; qvEx512* | P*rgef-1::sdn-1* | 69 | 35 | 5.7 |
| *sdn-1(zh20); arIs164; qvEx513* | P*rgef-1::sdn-1* | 74 | 41 | 5.5 |
| *sdn-1(zh20); arIs164; qvEx516* | P*rab-3::sdn-1* | 108 | 51 | 4.8 |
| *sdn-1(zh20); arIs164; qvEx474* | P*rab-3::sdn-1* | 103 | 38 | 4.8 |
| *sdn-1(zh20); arIs164; qvEx510* | P*mec-7::sdn-1* | 101 | 45 | 4.9 |
| *sdn-1(zh20); arIs164; qvEx439* | P*mec-7::sdn-1* | 108 | 54 | 4.8 |
| *sdn-1(zh20); arIs164; qvEx440* | P*mec-7::sdn-1* | 111 | 59 | 4.7 |
| *sdn-1(zh20); arIs164; qvEx514* | P*myo-3::sdn-1* | 100 | 43 | 5.0 |
| *sdn-1(zh20); arIs164; qvEx515* | P*myo-3::sdn-1* | 100 | 45 | 5.0 |
| *sdn-1(zh20); arIs164; qvEx465* | P*myo-3::sdn-1* | 101 | 42 | 4.9 |
| *sdn-1(zh20); arIs164; qvEx475* | P*myo-2::sdn-1* | 103 | 58 | 4.9 |
| *sdn-1(zh20); arIs164; qvEx517* | P*myo-2::sdn-1* | 140 | 52 | 4.2 |
| **Figure 4C** | | | | |

| | | | |
|---|---|---|---|
| *arIs164* | 339 | 0 | 0 |
| *unc-6(ev400); arIs164* | 355 | 13 | 1.8 |
| *unc-6(e78); arIs164* | 212 | 6 | 1.6 |
| *unc-40(e1430); arIs164* | 301 | 5 | 1.3 |
| *unc-40(e271); arIs164* | 208 | 7 | 1.8 |
| *unc-5(e53); arIs164* | 459 | 23 | 2.0 |
| *unc-40(e1430); unc-5(e53); arIs164* | 328 | 27 | 2.4 |
| **Figure 4D** (in grey when same data as in 4C) | | | |
| *arIs164* | 339 | 0 | 0 |
| *sdn-1(zh20); arIs164* | 102 | 49 | 4.9 |
| *unc-5(e53); arIs164* | 459 | 23 | 2.0 |
| *unc-5(e53); sdn-1(zh20); arIs164* | 123 | 56 | 4.5 |
| *unc-40(e1430); arIs164* | 301 | 5 | 1.3 |
| *unc-40(e1430); sdn-1(zh20); arIs164* | 141 | 56 | 4.4 |
| **Figure 5A** | | | |
| *arIs164* | 171 | 0 | 0 |
| *sdn-1(zh20); arIs164* | 111 | 48 | 4.8 |
| *ced-10(n1993); arIs164* | 248 | 2 | 0.8 |
| *ced-10(n1993); sdn-1(zh20); arIs164* | 209 | 20 | 2.8 |
| **Figure 5B** | | | |
| *arIs164* | 100 | 0 | 0 |
| *sdn-1(zh20); arIs164* | 124 | 47 | 4.5 |
| *mig-2(mu28); arIs164* | 146 | 3 | 1.5 |
| *sdn-1(zh20) mig-2(mu28); arIs164* | 158 | 17 | 3.0 |

N, number of animals in which the number of canals was examined. s.e.p., standard error of the proportion.

**S5 Table**. List of transgenic strains used.

| Strain | Genotype | Transgene | Reference |
|---|---|---|---|
| **Transgenic Lines** | | | |
| VQ1434 | *sdn-1(zh20) X; arIs164 V; qvEx437* | pCB420 (P*glt-3::sdn-1*) at 0.2ng/µl, P*unc-122::rfp*, pBSK+. Line #1 | This study |
| VQ1435 | *sdn-1(zh20) X; arIs164 V; qvEx438* | pCB420 (P*glt-3::sdn-1*) at 0.2ng/µl, P*unc-122::rfp*, pBSK+. Line #2 | This study |
| VQ1645 | *sdn-1(zh20) X; arIs164 V; qvEx509* | pCB420 (P*glt-3::sdn-1*) at 0.2ng/µl, P*unc-122::rfp*, pBSK+. Line #3 | This study |
| VQ1301 | *sdn-1(zh20) X; arIs164 V; qvEx352* | pCB265 (P*grd-10::sdn-1*) at 25ng/µl, P*unc-122::rfp*, pBSK+. Line #1 | This study |
| VQ1302 | *sdn-1(zh20) X; arIs164 V; qvEx353* | pCB265 (P*grd-10::sdn-1*) at 25ng/µl, P*unc-122::rfp*, pBSK+. Line #2 | This study |
| VQ1303 | *sdn-1(zh20) X; arIs164 V; qvEx354* | pCB265 (P*grd-10::sdn-1*) at 25ng/µl, P*unc-122::rfp*, pBSK+. Line #3 | This study |
| VQ1647 | *sdn-1(zh20) X; arIs164 V; qvEx511* | pCB425 (P*rgef-1::sdn-1*) at 25ng/µl, P*unc-122::rfp*, pBSK+. Line #1 | This study |
| VQ1648 | *sdn-1(zh20) X; arIs164 V; qvEx512* | pCB425 (P*rgef-1::sdn-1*) at 25ng/µl, P*unc-122::rfp*, pBSK+. Line #2 | This study |
| VQ1649 | *sdn-1(zh20) X; arIs164 V; qvEx513* | pCB425 (Prgef-*1::sdn-1*) at 25ng/µl, P*unc-122::rfp*, pBSK+. Line #3 | This study |
| VQ1652 | *sdn-1(zh20) X; arIs164 V; qvEx516* | pCB436 (P*rab-3::sdn-1*) ) at 5ng/µl, P*unc-122::rfp*, pBSK+. Line #1 | This study |
| VQ1564 | *sdn-1(zh20) X; arIs164 V; qvEx474* | pCB436 (P*rab-3::sdn-1*) at 5ng/µl, P*unc-122::rfp*, pBSK+. Line #2 | This study |
| VQ1646 | *sdn-1(zh20) X; arIs164 V; qvEx510* | pCB242 (P*mec-7::sdn-1*) at 25ng/µl, P*unc-122::rfp*, pBSK+. Line #1 | This study |
| VQ1446 | *sdn-1(zh20) X; arIs164 V; qvEx439* | pCB242 (P*mec-7::sdn-1*) at 25ng/µl, P*unc-122::rfp*, pBSK+. Line #2 | This study |
| VQ1447 | *sdn-1(zh20) X; arIs164 V; qvEx440* | pCB242 (P*mec-7::sdn-1*) at 25ng/µl, P*unc-122::rfp*, pBSK+. Line #3 | This study |
| VQ1650 | *sdn-1(zh20) X; arIs164 V; qvEx514* | pCB423 (P*myo-3::sdn-1*) at 10ng/µl, P*unc-122::rfp*, pBSK+. Line #1 | This study |
| VQ1651 | *sdn-1(zh20) X; arIs164 V; qvEx515* | pCB423 (P*myo-3::sdn-1*) ) at 10ng/µl, P*unc-122::rfp*, pBSK+. Line #2 | This study |
| VQ1533 | *sdn-1(zh20) X; arIs164 V; qvEx465* | pCB423 (P*myo-3::sdn-1*) ) at 10ng/µl, P*unc-122::rfp*, pBSK+. Line #3 | This study |
| VQ1565 | *sdn-1(zh20) X; arIs164 V; qvEx475* | pCB427 (P*myo-2::sdn-1*) at 1ng/µl, P*unc-122::rfp*, pBSK+. Line #1 | This study |
| VQ1653 | *sdn-1(zh20) X; arIs164 V; qvEx517* | pCB427 (P*myo-2::sdn-1*) at 1ng/µl, P*unc-122::rfp*, pBSK+. Line #2 | This study |

**Figure S1. Relative expression of the proteins of interest** from the Fig. 1 (A), Fig. 4 (B) and Fig. 5 (C) in the excretory cell during its development (270 min – 690 min of embryonic development). Data extracted from the viscello website on single-cell transcriptomic analysis during the embryonical development of C. *elegans*.

Ref citation viscello: Q. Zhu, J. I. Murray, K. Tan, J. Kim, qinzhu/VisCello: VisCello v1.0.0 (2019; https://zenodo.org/record/3262313)

### Figure S1

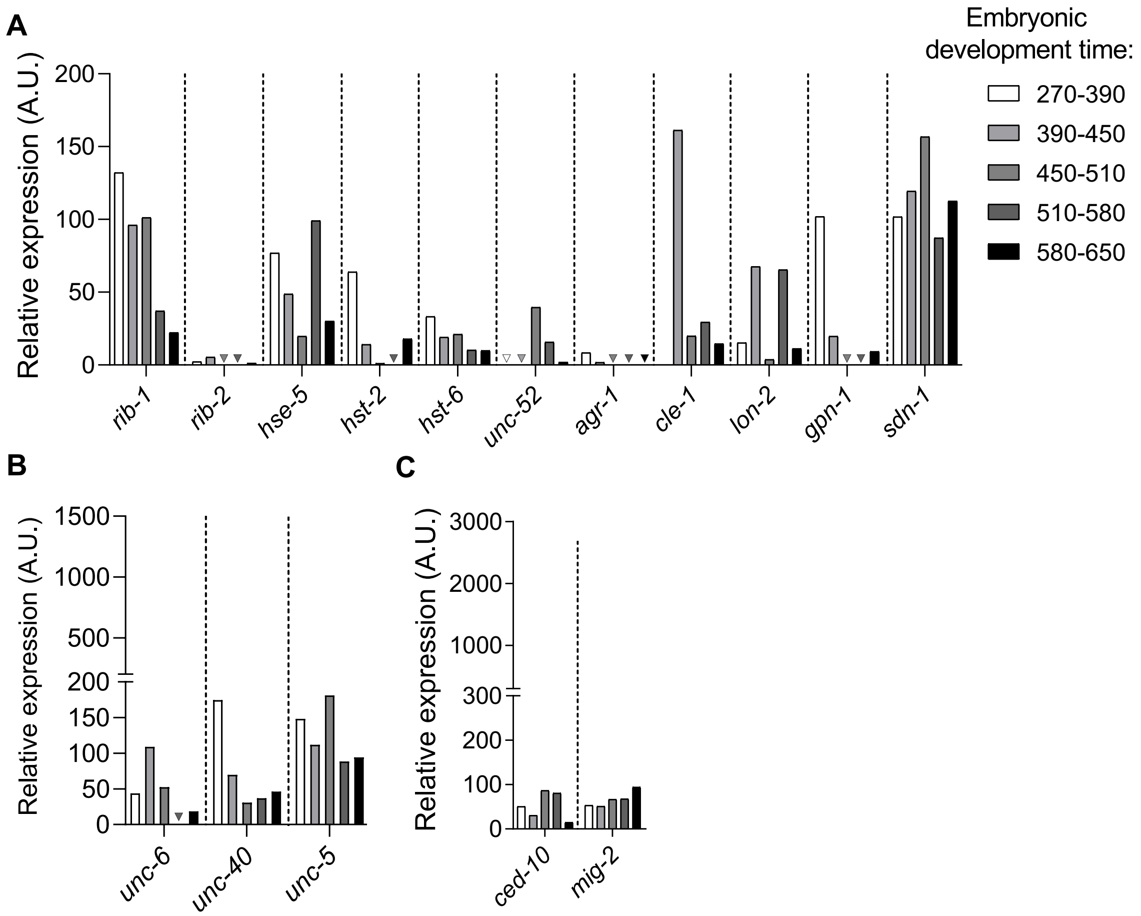

A
B
C
Relative expression (A.U.)
Embryonic development time:
270-390
390-450
450-510
510-580
580-650
rib-1
rib-2
hse-5
hst-2
hst-6
unc-52
agr-1
cle-1
lon-2
gpn-1
sdn-1
unc-6
unc-40
unc-5
ced-10
mig-2
